# Targeted Long-Read Bisulfite Sequencing for Promoter Methylation Analysis in Severe Preterm Birth

**DOI:** 10.1101/2024.03.04.583424

**Authors:** Silvana Pereyra, Angela Sardina, Rita Neumann, Celia May, Rossana Sapiro, Bernardo Bertoni, Mónica Cappetta

## Abstract

DNA methylation plays a critical role in the dynamics of gene expression regulation and the development of various disorders. Whole-genome bisulfite sequencing can provide single base resolution of CpG methylation levels and is the “gold standard” for DNA methylation quantification, but it also has a high cost. In contrast, targeted sequencing is optimal when focusing on specific candidate regions, while providing sufficient sequencing depth. Here, we present a targeted bisulfite sequencing approach to study the methylation status of regions of interest. We amplify selected regions from bisulfite-treated DNA and sequence them using Nanopore sequencing. In this work, we applied this workflow to candidate gene promoters for severe premature labor in a Latin American population.

We successfully amplified fragments over 1 Kb in length using long PCR conditions for 12 genes that were barcoded per sample and pooled to be sequenced on MinION flow cells. This approach achieved high sequencing depths, ensuring reliable DNAm estimation. We found significant hypomethylation of the *MIR155HG* gene promoter in severe preterm birth samples, which is concordant with reported gene expression changes.

We demonstrate that combining bisulfite DNA treatment with pooled long-read sequencing is a cost- and time-effective method to evaluate DNAm in several targeted regions and several samples in parallel. This study provides proof-of-concept for larger studies, demonstrating the applicability and high scalability of our assay to any locus of interest. Our experience suggests that this approach can be easily transferred to other diagnostic questions.

## 1 Introduction

Recent studies highlight the critical role of epigenetic modifications, including DNA methylation, in regulating gene expression and contributing to the development of various diseases. DNA methylation, the addition of a methyl group to the 5-prime position of a cytosine base in cytosine-phosphate-guanosine (CpG) dinucleotides, regulates gene expression by altering DNA accessibility to the transcriptional machinery in gene promoters and regulation sequences. CpG islands (CGI) are regions with a high density of CpGs, and methylation in CGI promoters plays an essential role in gene regulation and transcriptional repression (Goldberg et al. 2007). DNA methylation alters chromatin structure and affects other epigenetics marks, such as histone modification. DNA methylation is also implicated in repressing transposable elements, playing a fundamental role in the maintenance of genomic stability (Greenberg and Bourc’his 2019). DNA methylation is tightly regulated and affected by environmental factors; therefore, aberrant DNA methylation can lead to various human diseases (Ortiz-Barahona et al. 2022). In this context, the detection of differential methylation is crucial for understanding the causes of differential gene expression and, potentially, human disorders.

Several methods are available for DNA methylation quantification. Mostly, DNA methylation analysis has relied on microarrays that focus on mapping CpG-rich regions, gene promoters, and cis-regulatory elements. However, even while this method assays >97% of RefSeq, it only captures approximately 3% of DNA methylation sites in the human genome, with a median of 18 sites per gene (Noguera-Castells et al. 2023). This results in a substantial loss of valuable information, and it is therefore very likely that many of the precise locations where aberrant DNA methylation (DNAm) drives the onset of a particular disease are either not included in present analyses, or are only somewhat indexed by proximal sites. The current ‘gold standard’ technique is whole-genome bisulfite sequencing, which is an alternate, more thorough method including all 28 million CpGs after bisulfite DNA treatment. Nonetheless, it is expensive, as adequate coverage is necessary to achieve reliable quantification of DNA methylation. It is frequently difficult to obtain the necessary depth for particular sites, which makes accurate quantification difficult (Flynn et al. 2022).

To overcome this limitation, targeted sequencing becomes optimal when a candidate region is defined. This approach ensures the necessary sequencing depth in specific regions of interest, providing a more reliable and comprehensive assessment of DNA methylation. This method offers the potential for cost reduction and includes various techniques like bait capture, nanopore Cas9-targeted sequencing (nCATS), adaptive sampling, and bisulfite-treated DNA amplification (Gilpatrick et al. 2020; Loose et al. 2016; Simpson et al. 2017).

Reduced representation bisulfite sequencing (RRBS) (Gu et al. 2011) allows assessment of a fraction of CpG sites (1.5-2M) at low cost, but often results in uneven coverage and scans non-variable regions (Morselli et al. 2021). Capture techniques (e.g. CpGiant, TruSeqEpic) use second generation short read technology to characterize bisulfite treated targets (Tanić et al. 2022). Hybridization capture-based enrichment using biotinylated probes can be used to capture oligos recovers targeted regions, but these probes can be expensive (Kacmarczyk et al. 2018). Alternatively, Nanopore sequencing can directly detect DNA methylation without the need for additional sample processing, as required in sodium bisulfite procedures, and can examine DNA methylation patterns directly from genomic DNA (Gilpatrick et al. 2020). However, as DNAm marks are not preserved during PCR amplification, only native genomic DNA can be analyzed, requiring an additional step to transform it into a targeted assay (i.e. nCATS). Nevertheless, nCATS can be an expensive and labor-intensive method, especially in undeveloped countries, requires high amounts of DNA (3 to 5 µg) (Gombert et al. 2023) and does not achieve high enrichment (Sun et al. 2021).

Bisulfite sequencing remains the gold standard for methylation analysis. Bisulfite treatment results in highly fragmented DNA, which typically prevents amplification of fragments larger than 300 to 500 bp (Tusnády et al. 2005). Larger size ranges, up to 1500 bp, have been achieved but only by employing commercial kits (Yang et al. 2015). Even so, bisulfite-treated DNA long amplification has the potential to be a low-cost approach, enabling the analysis of numerous regions and scalability for cost-effective examination of a large number of samples. This approach proves ideal for population studies on complex diseases.

Preterm birth (PTB), as well as other complex diseases, is regulated by changes in DNA methylation (Park et al. 2020). It is defined as the delivery of an infant before 37 weeks of gestation and is the result of the interaction of genetic and environmental components, representing a multifactorial syndrome (Romero et al. 2014). PTB can be classified as severe (< 34 weeks of gestation) or moderate (34 to 37 weeks). Preterm birth can occur spontaneously or be induced by a physician through induction of labor or cesarean section, which may be medically indicated. Despite advances in understanding the risk factors and mechanisms underlying preterm birth, the underlying basis of PTB is still not fully understood, primarily because of the complex interplay of genetic, environmental, and host factors. Therefore, it is important to develop multiple approaches that integrate genetic, transcriptomic, epigenetic, and epidemiological data to identify biomarkers that will enable advances in translational medicine.

As with other diseases, most genetic research on preterm birth is conducted on people of European descent (Fatumo et al. 2022; Sirugo et al. 2019). The lack of ethnic diversity in studies of preterm birth means that published results are not necessarily generalizable to other populations, as distinct demographic histories can lead to unique genetic variants associated with a given phenotype (Martin et al. 2020). Thus, exploring genetic diversity in underrepresented populations is crucial for a comprehensive understanding of health and disease in diverse human populations. To this end, our group has made efforts in integrated genomics and epigenetics studies on complex diseases such as cancer and preterm birth, aiming to identify biomarkers in Latin American populations (Brignoni et al. 2020; Cappetta et al. 2015, 2021; Pereyra et al. 2016, 2019).

In this study, we have developed an analysis pipeline to investigate the methylation status of multiple gene promoters. We tested this workflow on candidate genes for spontaneous severe preterm labor in a Uruguayan population. We selected genes previously found to be differentially expressed in chorioamniotic tissue during preterm labor (Pereyra et al. 2019). Our results demonstrate the use of a cost-effective approach to analyze DNA methylation in targeted regions.

## 2 RESULTS

A total of 6 chorioamniotic tissue samples from term newborns and 5 from severe preterm newborns were analyzed. A subset of the epidemiologic variables analyzed are shown in Supplementary Table S1. There were no significant differences among groups in the sex of the infants, maternal age, number of previous gestations, premature rupture of membranes (PROM) or anemia (p > 0.05). Preterm infants had significantly lower birth weights and earlier gestational ages at delivery. None of the mothers actively smoked cigarettes during pregnancy in this study, and the proportion of passive smoking showed no significant difference between PTB and control. No mothers had previously had preterm births, nor presented hypertension, preeclampsia, intrauterine growth restriction (IUGR) or hemorrhages during pregnancy.

Four Oxford Nanopore sequencing Minion runs were performed. After alignment and filtering reads with quality scores greater than 7, a total of 60.914 reads were mapped to targeted genes. After filtering per coverage (>30 reads per site), a total of 788 unique CpG sites were available for analyses in the targeted genes. These CpG are distributed among promoter genes as shown in Table 1. The average sequencing depth per CpG was 619x, while the mean coverage per sequencing run varied between 325x and 834x. Taken all data together, mean coverage per site and gene ranged from 61x to 1182x (Table 1). Due to different PCR efficiencies, sequencing data were not obtained for all genes for all samples.

**Table 1.** Chromosomal and relative positions of amplified regions. *Logarithmic Fold Change value as reported in Pereyra et al. (2019), reflecting differences in gene expression level between severe preterm birth and term birth newborns.

| Gene | Chromosomal position of amplified region | Relative position to TSS | CpG context | Number of analyzed | Mean Coverage | LogFC value* |
| --- | --- | --- | --- | --- | --- | --- |

|  |  |  |  | CpGs |  |  |
| --- | --- | --- | --- | --- | --- | --- |
| *ACCS* | chr11:44065877-4406683<br>9 | (-392) - (+570) | Island | 72 | 81.24 | -2.79 |
| *ANKRD24* | chr19:4198696-4199647 | (+607) -<br>(+1558) | shore | 22 | 284.01 | -2.21 |
| *BIRC3* | chr11:102317048-102317<br>759 | (-435) - (+276) | Island | 39 | 151.23 | 3.36 |
| *CXCL2* | chr4:74098538-74099133 | (+62) - (+657) | Island | 93 | 701.37 | 4.96 |
| *EGR3* | chr8:22694093-22695046 | (-2682) -<br>(-1729) | Island | 65 | 61.68 | 3.67 |
| *GK* | chrX:30652839-30653338 | (-583) - (-84) | Island | 54 | 1000.89 | 5.04 |
| *MAMDC2* | chr9:70043447-70044211 | (-133) - (+631) | Island | 79 | 1182.61 | -2.23 |
| *MIR155H*<br>*G* | chr21:25562022-2556261<br>9 | (-122) - (+475) | Island | 60 | 302.17 | 2.66 |
| *NRN1* | chr6:6003310-6004084 | (-59) - (+715) | Island | 138 | 612.79 | -2.19 |
| *PIK3AP1* | chr10:96720099-9672091<br>3 | (+415) - (-399) | Island | 65 | 68.73 | 4.37 |
| *STEAP1* | chr7:90153483-90154614 | (-985) - (+146) | Island | 84 | 431.86 | 4.17 |
| *WNT1* | chr12:48976387-4897755<br>7 | (-1934) - (-764) | shore | 17 | 87.14 | -3.29 |

Hierarchical clustering of sample methylation profiles using either Pearson’s correlation distance or principal component analysis did not group samples based on their categories. (Supplementary Figure S1). Methylation percentage levels per CpG per gene showed high variability across gene promoters (Supplementary Figure S2). In spite of the low sample number, a differentially methylated region (DMR) analysis was performed and we detected one differentially methylated region between preterms and controls in *CXCL2* gene, chr4:74099080-74099111, comprising 3 CpGs (FDR < 0.05).

When CpGs were grouped per gene and per condition, significant methylation differences were detected between cases and controls in 7 gene promoters: *ANKRD24*, *MIR155HG*, *CXCL2*, *NRN1*, *STEAP1*, *MAMDC2* and *GK* (Wilcoxon rank sum test, adjusted FDR p value < 0.05, Figure 1, Supplementary Table S2). Changes in methylation observed here (Figure 1) were compared with RNA expression levels and found to be concordant for *MIR155HG* gene, but not for the remaining analyzed genes (Table 1). In the *MIR155HG* promoter, we detected hypomethylation in severe preterm births, while mRNA levels were previously reported to be overexpressed in this group (Pereyra et al. 2019) (Figure 2).

**Figure 1.**
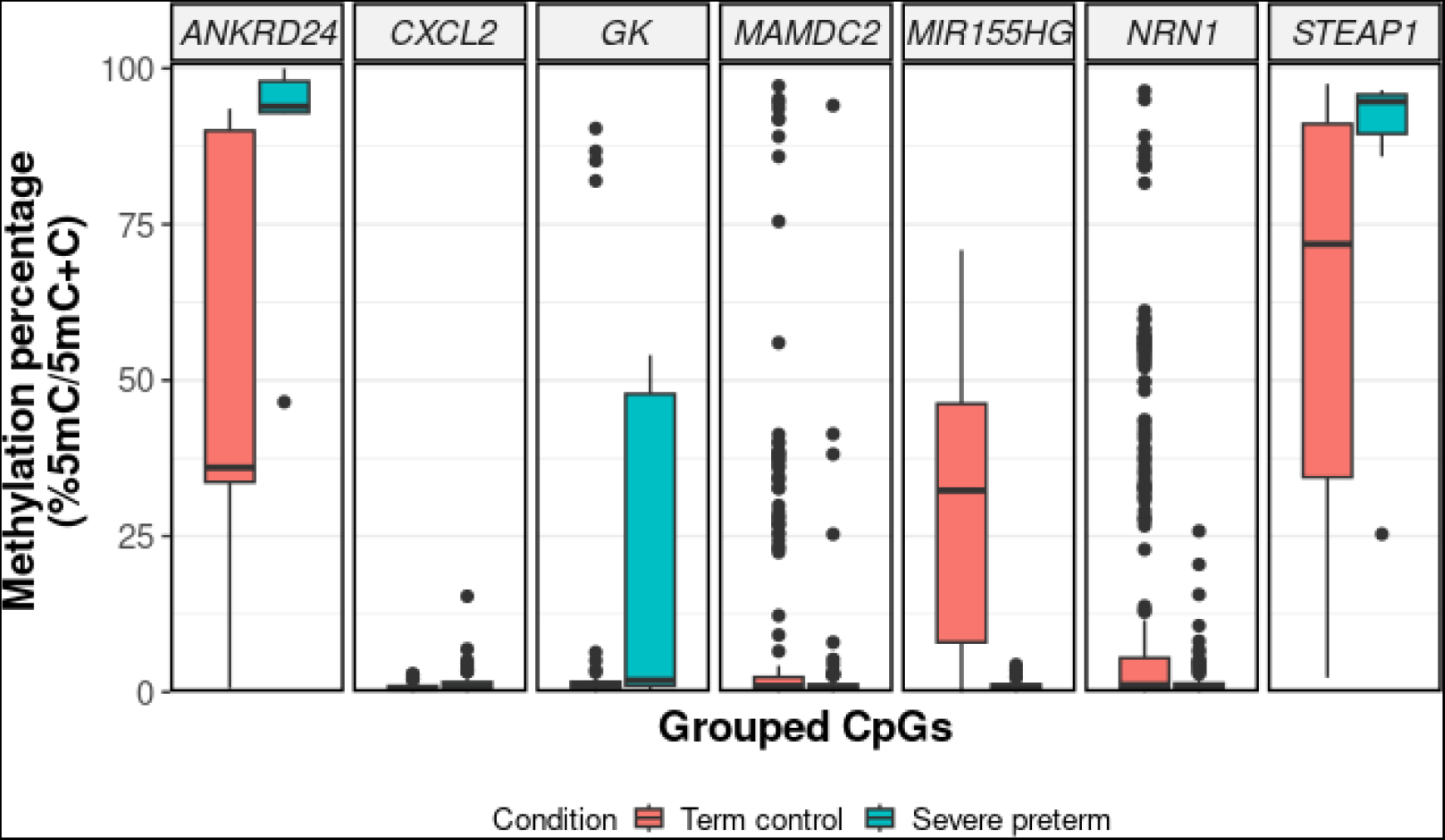
DNA methylation level of grouped CpG sites per gene which showed significant differences in chorioamniotic membranes between severe preterm and term patients. The boxes represent the interquartile ranges, and the lines across the boxes indicate the median value. Statistically significant differences between severe preterm and term patients were determined using Wilcoxon rank sum test and adjusted by FDR (p < 0.05).

**Figure 2.**
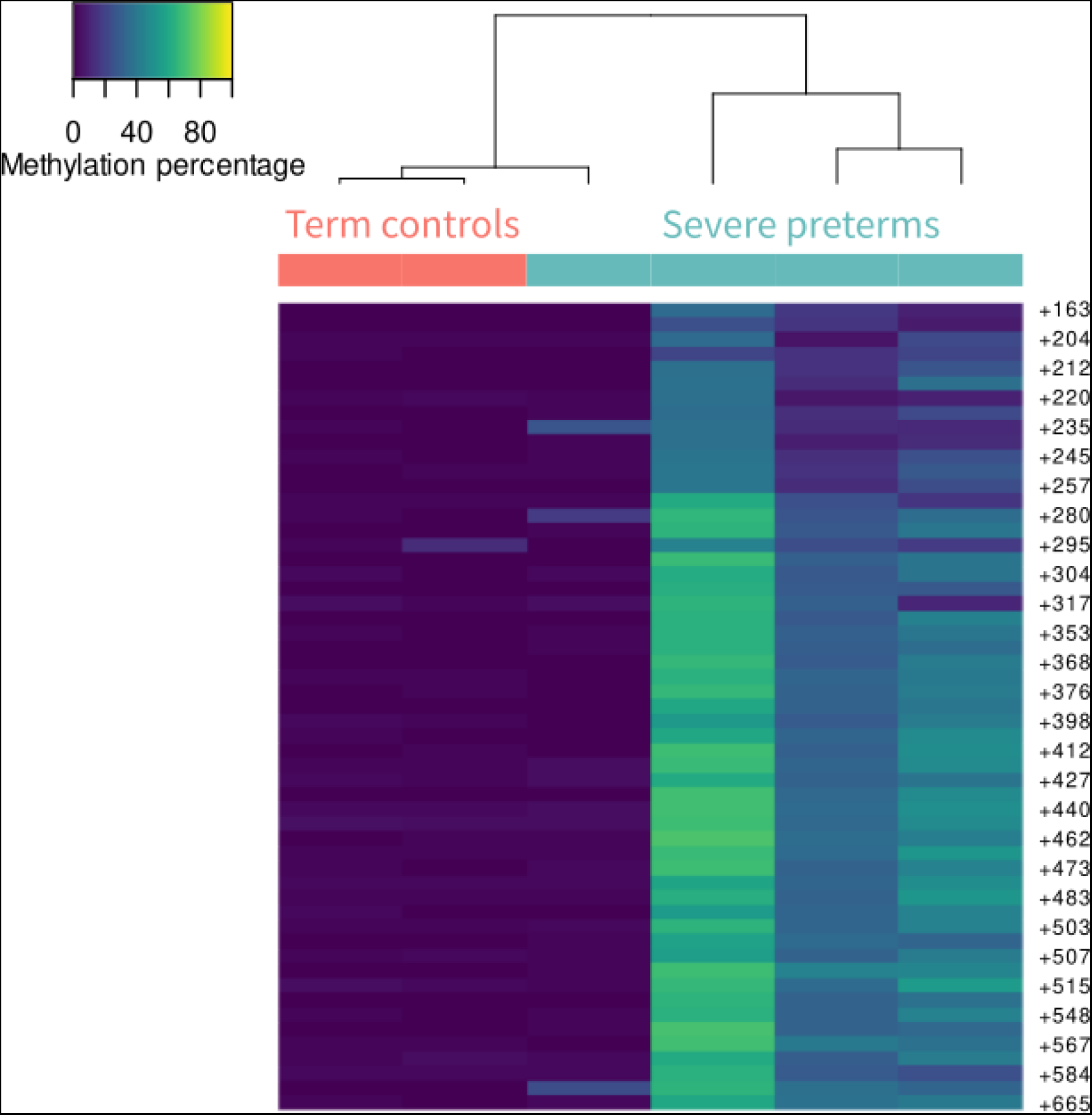
Methylation percentage per CpG site in term controls and severe preterms for *MIR155HG.* Each row represents a different CpG site, while columns represent individual samples. Unsupervised hierarchical clustering with complete linkage and Euclidean distance was used to group samples based on their methylation profiles. CpG sites are annotated relative to the transcription start sites (TSS). Sequencing data were not obtained for all samples.

In order to assess if combining methylation and gene expression data can discriminate between cases and controls, a combined reduction dimension analysis was performed. Initially, MDS analysis of methylation data did not distinctly cluster samples based on their condition (Figure 3A). However, control samples are more concentrated along the first dimension axis, while severe preterm samples are positioned along the second dimension axis, suggesting a differential methylation pattern. Upon integrating both methylation and transcriptome data for samples where both datasets were available, severe preterm samples clustered together, while control samples were dispersed (Figure 3B).

**Figure 3.**
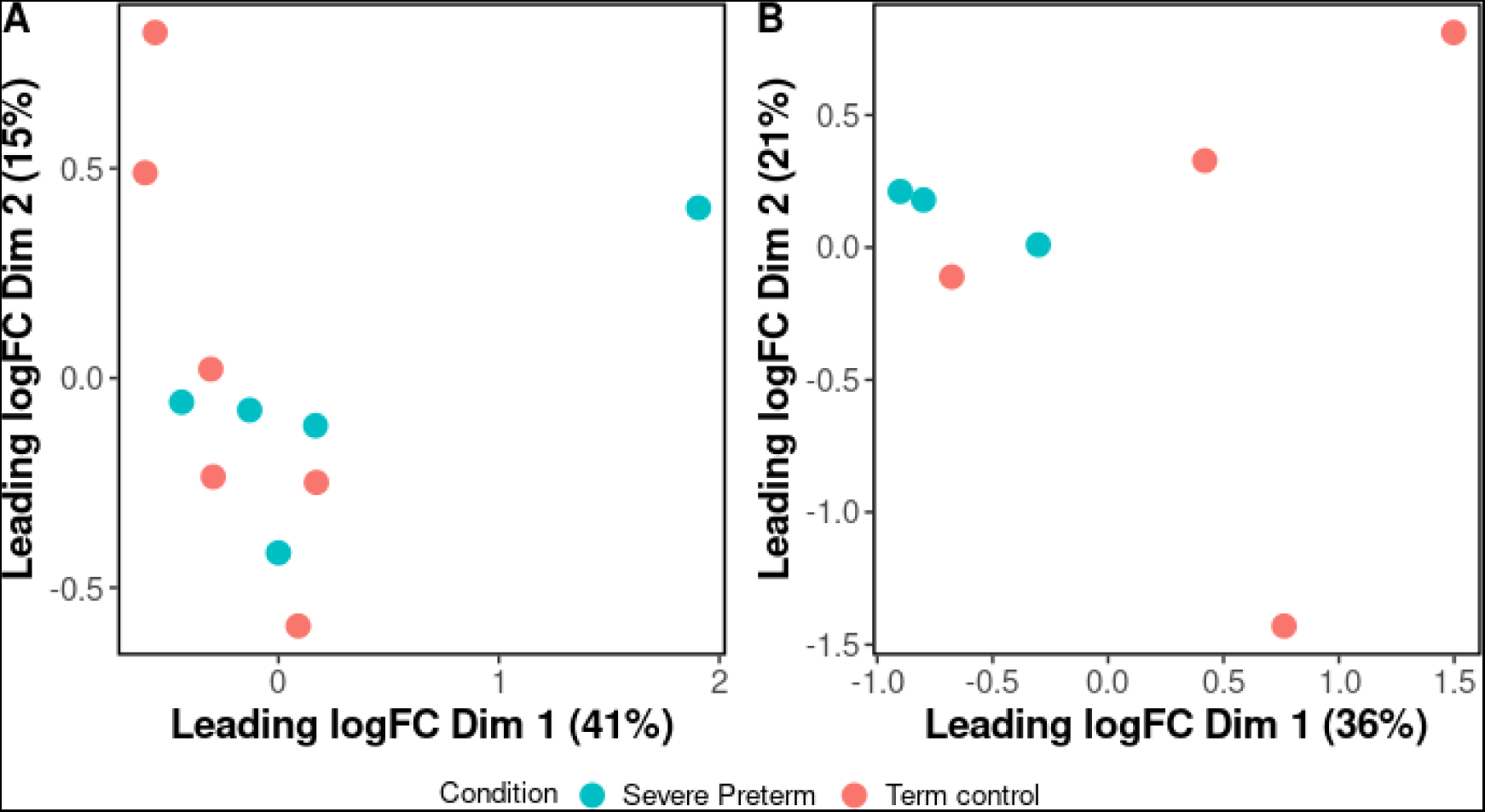
**(A)** Multidimensional scaling (MDS) plot generated based on the matrix of methylation levels and Euclidean distances for all samples. **(B)** MDS plot of methylation and transcriptome data for 7 samples.

## 3 DISCUSSION

In this study, we report an accurate and cost- and time-effective method to characterize promoter DNA methylation using nanopore sequencing. PCR amplicons derived from bisulfite-treated DNA samples from severe preterm and term newborns were pooled and sequenced on MinION flow cells. Custom-designed primers were used to target selected regions of 12 candidate gene promoters.

The targeted bisulfite sequencing approach presented offers several advantages over alternative targeted approaches. Notably, our approach achieves exceptionally high coverage per gene, reaching up to 1100x in total and up to 834x per run. Read depth has a crucial effect on the accuracy of DNAm estimates, particularly at sites with intermediate DNAm levels, which may be erroneously classified as methylated or unmethylated at low read depths (Seiler Vellame et al. 2021). Masser et al. (2013) identified a 1000x read depth is necessary for reliable methylation estimates. Our approach ensures sufficient depth at each site in all samples, ensuring reliable DNAm estimation. Our approach therefore performs better than nCATS assays, which often yield much lower coverages with considerable variance, ranging from means of 350x to as low as <10x (Skowronek et al. 2022; Wieting et al. 2023; Alfano et al. 2022; Iyer et al. 2022).

Generally, because bisulfite-treated DNA has a high amount of uracil homopolymers, it is often difficult to amplify, yielding a low typical bisulfite PCR size range (∼300-500 bp) (Yang et al. 2015). However, the use of long PCR conditions allowed us to amplify fragments of over 1 Kb in length successfully.

The approach presented here allows the analysis of DNAm across multiple candidate regions and it is highly customizable and scalable. The number of amplicons to be sequenced can be comfortably increased without compromising performance. It is also possible to increase the number of barcoded samples run on each flow cell. It’s important to note that in our experiment, each flow cell is underutilized, allowing to potentially scale up the assay to optimize resources efficiently. While we used MinION flow cells, capable of sequencing up to 48 Gb, this methodology could be adapted to Flongle flow cells, which have a yield of up to 2.6 Gb, further reducing costs. This makes our approach ideal for research laboratories analyzing DNAm with limited resources, a common scenario in geographic regions with underrepresented populations in genomic studies. Facilitating access to research methods for studying genetic and epigenetic variation in underrepresented populations is crucial; without data from diverse populations, our understanding of genomic influences on health would be incomplete, potentially exacerbating health disparities and inequities.

The customizability of our approach means that, unlike microarrays, which are primarily designed for model species, this technique can also be adapted for non-model species with little effort. Alternatively, adaptive sampling is a PCR-free method for analyzing DNAm directly from native DNA in targeted regions (Kovaka et al. 2021; Payne et al. 2021). Adaptive sampling does not require treating samples with sodium bisulfite, nor designing or running PCRs. However, it is currently only optimized for MinION flow cells, increasing assay costs unless run at scale.

Epigenetic regulation has proved to be a relevant mechanism for PTB in regulatory tissues (Park et al. 2020), but no studies have been performed in chorioamniotic membranes to date. DNA methylation differences between term and preterm births have been reported in fetal amnios (Parets et al. 2013; Kim et al. 2013), placenta (Schoorlemmer et al. 2020), maternal blood at birth (You et al. 2021), umbilical cord and blood (Wu et al. 2019; Wang et al. 2019). To the best of our knowledge, this work marks the first targeted long reads sequencing effort using Nanopore technology to study a Latin American population, and the first one to analyze DNAm data from chorioamniotic membranes for PTB. We report methylation levels for samples of chorioamniotic membranes and contrast the methylation differences found with existing transcriptome data in the same samples. *MIR155HG* is the only gene for which the direction of DNA methylation changes in its promoter are concordant with reported gene expression changes. *MIR155HG* encodes a precursor RNA of microRNA-155 (miRNA-155) which has been identified as a regulator of inflammatory responses in several pathologies (Mahesh and Biswas 2019; Kurowska-Stolarska et al. 2011; Peng et al. 2019). Aberrant miR-155-5p overexpression can be involved in preeclampsia pathogenesis, and its detection in maternal serum samples obtained during gestation was one of the best predictive biomarkers of preeclampsia, even before the onset of symptoms (Srinivasan et al. 2020). In our data, we found the analyzed region of the *MIR155HG* gene promoter to be hypomethylated in preterm birth, which may be highly associated with high *MIR155HG* gene expression in preterm birth in chorioamniotic membranes. The analyzed region includes the most proximal CpG island to the transcription start site. *MIR155HG* was also found hypomethylated in glioma patients, and showed a similar correlation to its elevated gene expression levels (Wu et al. 2022). This highlights the importance of epigenetic regulation in inflammatory pathologies like PTB and cancer.

Lack of concordance for the remaining analyzed genes may be due to the presence of different gene regulation mechanisms to DNAm, such as non-coding RNA regulation, histone modification or chromatin remodeling. For example, *PIK3AP1* has an aberrant expression in several cancers, where *PIK3AP1* expression in cancer cells is silenced by overexpression of miRNAs (Guo et al. 2021; Li et al. 2022). Also, our design only analyzed a limited promoter region per gene (1Kb approximately, especially targeting CpG islands), which may not necessarily represent the DNAm levels of the whole gene promoter. In the future, a larger region should be analyzed to detect changes, as DNAm along the promoter region may not be homogeneous (Irizarry et al. 2009).

Despite the discordance observed between promoter methylation and gene expression data, MDS analysis integrating both methylation and transcriptome data for samples was able to cluster severe PTB cases. This integrated approach demonstrates potential for future sample categorization in this disease.

Limitations of this study include the small number of samples due to the pilot nature of the study, which particularly limits the ability to detect differential methylation between groups. Nevertheless, this study provides a proof-of-concept for the design of larger studies needed to confirm the observed methylation trends.

We demonstrate that combining bisulfite DNA treatment with pooled long-read sequencing is a reliable way to evaluate DNAm in several targeted regions and several samples in parallel. We developed this assay for 12 promoter regions relevant to preterm birth, but the method is applicable and highly scalable to any locus of interest. Our experience suggests that this approach can be easily transferred to other diagnostic questions.

## 4 METHODS

### 4.1 Study population

Controls and cases were term and spontaneous severe preterm deliveries respectively from unrelated offspring of women receiving obstetrical care at the Centro Hospitalario Pereira Rossell, Montevideo, Uruguay, recruited in a previous study described in Pereyra et al. (2019). Briefly, severe preterm chorioamnion tissues were collected immediately after labor from pregnancies complicated by birth before 33 weeks of gestational age (GA). Term chorioamnion tissues were obtained from uncomplicated pregnancies delivered after 37 weeks of GA.

The ethics committee of the Facultad de Medicina of the Universidad de la República (Uruguay) and the Ethics committee of the Centro Hospitalario Pereira Rossell (Uruguay) both approved the study protocol (Facultad de Medicina Board resolution N° 071140-000907-11). Written informed consent was obtained from mothers prior to the collection of biological material.

Maternal demographic and patronymic data were collected through questionnaires filled out by the mothers after delivery. Clinical and obstetric data were obtained from the Perinatal Information System, which consists of basic perinatal clinical records developed by the Latin American Center for Perinatology (CLAP) from WHO/PAHO (De Mucio et al. 2016). A subset of epidemiological variables collected from the participants was examined to assess potential differences among groups. Gestational age, newborn weight, maternal age and previous gestations were analyzed as continuous variables and were assessed using Student’s t-test, while newborn sex, PROM, IUGR, preeclampsia, anemia, hypertension and smoking were considered qualitative variables and were assessed using Chi-square tests. All statistical analyses were carried out using R programming language (http://www.r-project.org/). Statistical significance was set at p < 0.05.

### 4.2 Bisulfite-treated DNA sequencing

DNA was extracted from chorioamniotic membranes using standardized salting-out methods. Five hundred nanograms of DNA per sample were bisulfite-treated using the Zymo EZ-96 DNA methylation kit (Zymo Research, Orange, CA, USA). A bisulfite sequencing PCR approach was applied to the promoters of 12 candidate genes (Supplementary Table S3). Candidate genes were selected based on the highest absolute logarithmic fold change values reported by Pereyra et al. (2019).

For each gene, a promoter region was selected to be amplified and analyzed. When possible, a CpG Island was included in the selected region. To optimize amplification of bisulfite treated DNA, long and nested PCR techniques were employed to amplify fragments of approximately 1Kb of length. Oligonucleotides for gene promoters were designed in Methyl Primer Express™ Software v1.0 (Applied Biosystems, USA), following directions in Wojdacz et al. (2008). Universal tail sequences as provided by Oxford Nanopore Technologies (forward primer: 5’-TTTCTGTTGGTGCTGATATTGC-3’, reverse primer: 5’-ACTTGCCTGTCGCTCTATCTTC-3’) were added to second PCR round primers, at their 5’ end. Designed oligonucleotides were checked on BiSearch Web Server (Tusnády et al. 2005). In total, the targeted regions cover ∼10Kb.

Amplification was performed using an Applied Biosystems Veriti™ Thermal Cycler (Thermo Fisher Scientific, Waltham, MA, USA) with the following PCR conditions for first PCR Round: 1 cycle of 96 °C for 5 seconds, gene-specific annealing temperature (Supplementary Table S3) for 1 minute and 64°C for 4 minutes, followed by 35 cycles of 95°C for 20 seconds, gene-specific annealing temperature for 30 seconds, and 64°C for 2 minutes. The second PCR round was incubated 96°C for 1 minute, and 32 cycles of 96°C for 20 seconds, gene-specific annealing temperature for 30 seconds, and 64°C for 1:30 minutes. First PCR rounds were performed in 10 μl reaction volumes, containing 1X in-house PCR buffer (Kauppi et al. 2009), 12 mM Tris, 0.3 μM of each Forward and Reverse gene primer, 1.2 units of HighTaq DNA Polymerase (Bioron, Germany), and 0.06 units of Velocity DNA Polymerase (Bioline, UK) and 50 ng of template DNA. Second PCR rounds were performed in 20 μl reaction volumes, containing 1X in-house PCR buffer, 12.5 mM Tris, 0.4 μM of each Forward and Reverse gene primer, 2.4 units of HighTaq DNA Polymerase (Bioron, Germany), and 0.12 units of Velocity DNA Polymerase (Bioline, UK) and 0.5 μL of first round PCR product as template.

### 4.3 Experimental design to optimize MinION flow cell use

To optimize MinION flow cell utilization, we concurrently sequenced 10 samples from our preterm birth study and 12 samples from a separate pathology study in each run. These studies investigate distinct gene promoters, with 12 genes analyzed in the preterm birth study and 15 in the other. Here, we exclusively present results from our preterm study.

For Nanopore sequencing, equal quantities of each amplicon per sample were pooled together. Libraries were prepared with a total of 300 ng of each pooled barcoded amplicons using a PCR Barcoding kit (SQK-PBK004) according to the manufacturer’s instructions. All sample libraries were multiplexed on a MinION device using a MinION flow cell (FLO-MIN106, R9.4.1, Oxford Nanopore), using MinKnow v21.06.13 software. Samples were sequenced for a total of 20 hours.

Raw FAST5 files produced by MinION were basecalled under high accuracy mode using the ONT basecaller Guppy v5.0.16 (Oxford Nanopore Technologies). Raw sequences were pre-processed to remove barcodes in MinKnow. A minimum quality score cut off of 7 was used for filtering low quality reads, employing Nanofilt (De Coster et al. 2018). The trimmed files were then aligned to the human reference genome (GRCh38) using Bismark v0.16.3 (Krueger and Andrews 2011), which was set to interpret only methylation in a ‘CG’ context, with the following parameters, optimized for long sequenced reads: --local --score_min G,1,5.4. Reads were visualized with the Integrative Genomics Viewer (IGV) browser (Thorvaldsdóttir et al. 2013). Aligned reads were further processed with Bismark Methylation Extractor (default parameters) to yield Bismark coverage files.

### 4.4 CpG site methylation analyses

All further analyses were carried out using the R programming language (http://www.r-project.org/). We filtered CpG sites retaining those with more than 30 reads, as reported in the Bismark coverage files. To remove spurious reads from library preparation and potential mapping artifacts, we used *Genomic Ranges* R package (Lawrence et al. 2013) to filter reads based on their genomic position for all selected genes. Wilcoxon rank sum test was performed to identify differentially methylated CpG sites between term and severe preterm groups, either individually or grouped. The p-values were adjusted for multiple testing using false discovery rate estimation (FDR), with a threshold p-value of < 0.05. For data exploration, we performed Pearson’s correlation analysis and principal component analysis on sites that were covered in all samples at 30× or greater, using the Methylkit package (Akalin et al. 2012) implemented in R. Cluster analysis of the differential methylated CpG sites was performed using unsupervised hierarchical clustering with complete linkage and Euclidean distance as a measure of similarity between samples.

A single-linkage clustering algorithm was implemented in the R/Bioconductor package BiSeq (Hebestreit and Klein 2015) to detect differentially methylated regions (DMRs). The data were reduced to CpG sites covered in 50% of the samples, comprising at least 7 CpG sites close enough to each other (max 100 bp). To reduce bias due to outlier coverages, the coverage of CpG sites was limited to the 90% quantile. Differently methylated clusters were detected with a false discovery rate (FDR) of 0.05. Within differentially methylated clusters, differentially methylated CpG sites were detected with an FDR of 0.05. Differentially methylated CpG sites were grouped in a DMR when they are less than 100 bp away from each other.

### 4.5 Combined reduction dimension analysis

A multidimensional scaling (MDS) analysis was employed as a dimension reduction technique to explore the dataset for patterns in the methylation values for the 11 samples assayed, with R package edgeR (Robinson et al. 2010). Additionally, when possible, methylation and expression levels within the same sample were compared. Since seven of the study’s examined samples had previously undergone RNAseq analysis (SRP139931) (Pereyra et al., 2019), it was possible to combine methylation and RNAseq data into an MDS, for the 12 genes analyzed in the present study.

## 5 DATA ACCESS

All raw and processed sequencing data generated in this study have been submitted to the NCBI SRA (https://www.ncbi.nlm.nih.gov/geo/) under accession number PRJNA1081111.

## 7 AUTHOR CONTRIBUTIONS

SP and MC conceived and designed the project. SP, MC, AS, and BB performed epigenetic experiments, and analyzed and interpreted the experimental data. CM and RN contributed to the development of long PCR experimental protocols. RS and BB contributed in the statistical analysis and provided feedback on the manuscript; and SP and MC wrote the manuscript. All authors read and approved the manuscript.

## 8 ACKNOWLEDGMENTS

The authors are indebted to participating mothers, whose generosity and cooperation have made this study possible. This study was partly supported by Programa de Desarrollo de las Ciencias Básicas (PEDECIBA) and Universidad de la República (UDELAR) to SP, MC and BB. MC, BB and RS are supported by Sistema Nacional de Investigadores, Agencia Nacional de Investigación e Innovación (ANII, Uruguay).

